# Neocortical pyramidal neurons with axons emerging from dendrites are frequent in non-primates, but rare in monkey and human

**DOI:** 10.1101/2021.12.24.474100

**Authors:** Petra Wahle, Eric Sobierajski, Ina Gasterstädt, Nadja Lehmann, Susanna Weber, Joachim H.R. Lübke, Maren Engelhardt, Claudia Distler, Gundela Meyer

## Abstract

The canonical view of neuronal function is that inputs are received by dendrites and somata, become integrated in the somatodendritic compartment and upon reaching a sufficient threshold, generate axonal output with axons emerging from the cell body. The latter is not necessarily the case. Instead, axons may originate from dendrites. The terms “axon carrying dendrite” (AcD) and “AcD neurons” have been coined to describe this feature. Here, we report on the diversity of axon origins in neocortical pyramidal cells. We found that in non-primates (rodent, cat, ferret, pig), 10-21% of pyramidal cells of layers II-VI had an AcD. In marked contrast, in macaque and human, this proportion was lower, and it was particularly low for supragranular neurons. Unexpectedly, pyramidal cells in the white matter of postnatal cat and aged human cortex exhibit AcDs to much higher percentages. In rodent hippocampus, AcD cells are functionally ‘privileged’, since inputs here can circumvent somatic integration and lead to immediate action potential initiation in the axon. Our findings expand the current knowledge regarding the distribution and proportion of AcD cells in neocortial regions of non-primate taxa, which strikingly differs from primates where these cells are mainly found in deeper layers and white matter.

## Introduction

The prevailing concept of neocortical pyramidal cell function proposes that excitatory inputs arrive via the dendrites, are integrated in the somatodendritic compartment, and upon reaching sufficient threshold, the axonal domain generates an action potential. The axon usually originates from the ventral aspect of the soma, starting with a short axon hillock followed by the axon initial segment (AIS), the electrogenic domain generating the action potential (reviewed by Kole and Brette, 2018). Already Ramon y Cajal suggested that impulses may bypass the soma and flow directly to the axon (reviewed by Triarhou, 2014). AcD are common in cortical inhibitory interneurons (Meyer, 1987; Wahle and Meyer, 1987; Meyer and Wahle, 1988; Höfflin et al., 2017). Further, upright, inverted and fusiform pyramidal neurons of supra – and infragranular layers display AcDs in Golgi impregnated or dye-injected cortex from rodents, lagomorphs, ungulates and carnivores (Smit and Uylings, 1975; van der Loos, 1976; Peters and Kara, 185; Ferrer et al., 1986a; Ferrer et al., 1986b; Hübener et al., 1990; Reblet et al., 1992; Matsubara et al., 1996; Prieto and Winer, 1999; Mendizabal-Zubiaga et al., 2007; Hamada et al., 2016; Ernst et al., 2018). In mouse hippocampal CA1 pyramidal cells, axons frequently emerge from basal dendrites (Thome et al., 2014). Multiphoton glutamate uncaging and patch clamp recordings revealed that input to the AcD is more efficient in eliciting an action potential than input onto regular dendrites (non-AcDs). Further, AcDs are intrinsically more excitable, generating dendritic spikes with higher probability and greater strength. Synaptic input onto AcDs generates action potentials with lower thresholds compared to non-AcDs, presumably due to the short electrotonic distance between input and the AIS. The anatomical diversity of axon origin plus the diversity of length and position of the AIS substantially impact the electrical behavior of neurons (reviewed by Kole and Brette, 2018). This begs the questions, how frequent AcD neurons are among the mammalian species, and whether AcD neurons also exist in primates. Our data suggest remarkable differences between phylogenetic orders and position in gray and white matter.

**Figure 1.**
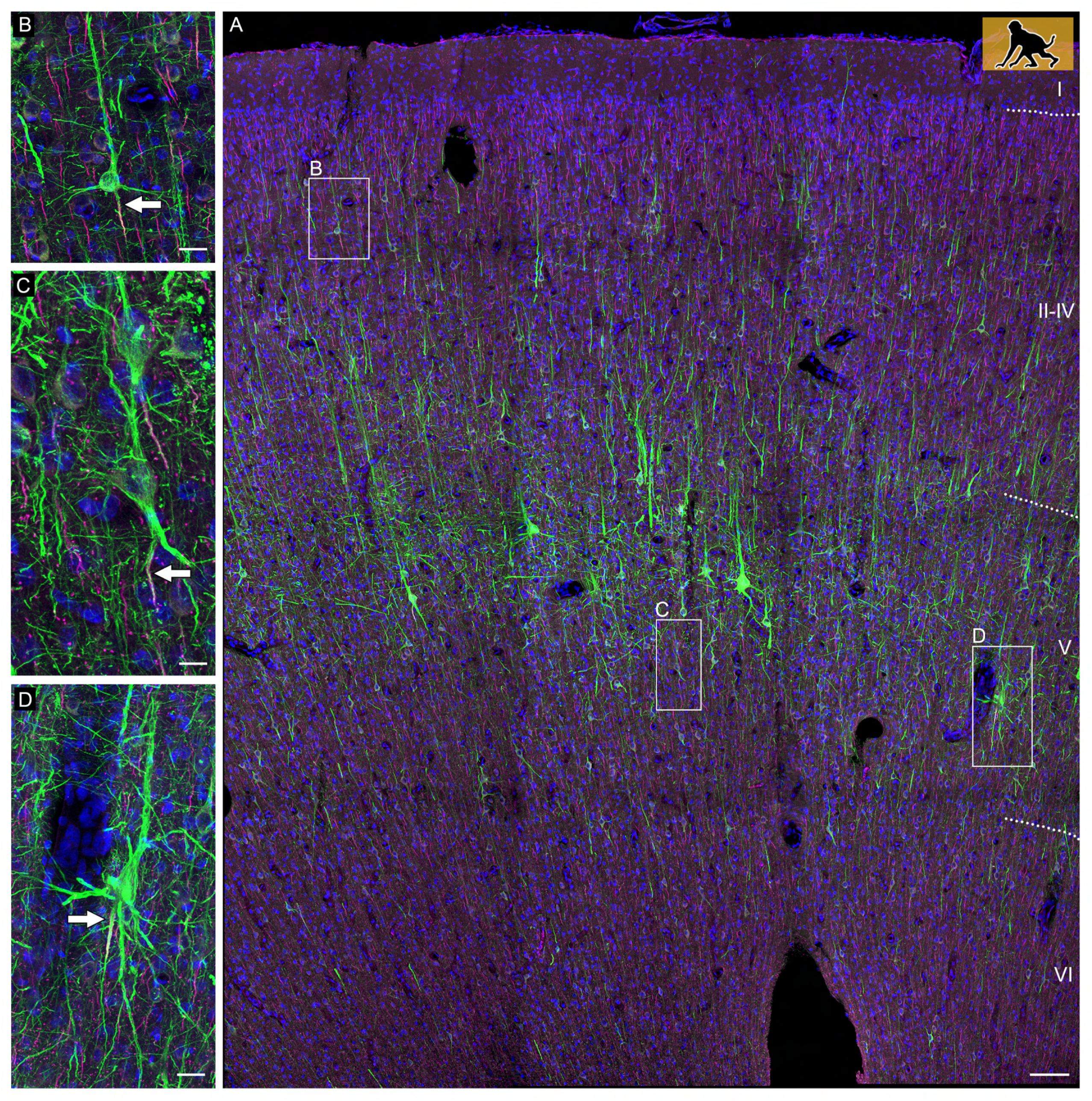
Confocal tile scan of dorsal neocortex (motor-premotor area) of P60 infant macaque. (A) Pyramidal cells were stained with SMI-32/βIV-spectrin to label dendrites and the AIS, respectively. Insets depict neurons with axons emerging from somata (B) or from an AcD (C), or a shared root (C). Axons indicated by arrows. Scale bars 100 μm for the tile scan, and 25 μm for the insets.

## Results

### AcD cells in adult cortex

***Figure 1A*** documents the diversity of axon origins in a P60 infant macaque monkey cortex with an axon originating from a soma (inset B), an AcD (inset B), and the “shared root” configuration (inset D), which we scored as somatic. Generally, AcDs were basal dendrites. AcD neurons of other species are shown in ***Figure 2A-E*** and ***Figure 2-figure supplement movie 1***.

**Figure 2.**
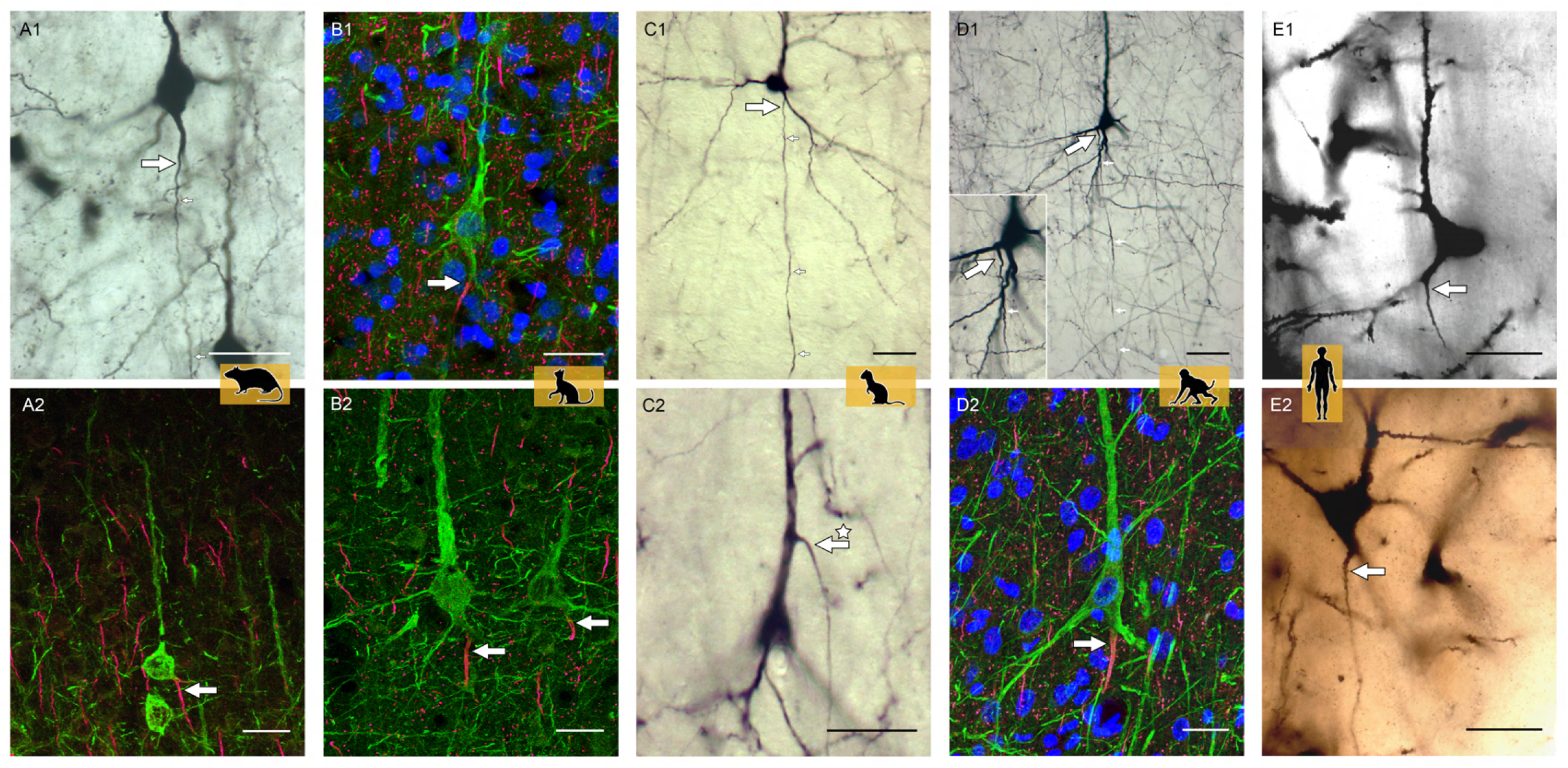
Representative AcD neurons. (A1, A2) from rat (biocytin, immunofluorescence); (B1, B2) cat (immunofluorescence); (C1, C2) ferret (biocytin); (D1, D2) monkey (biocytin immunofluorescence), the inset shows the axon origin at higher magnification; (E1, E2) human (Golgi method; D2 is a montage of two fotos). Apical AcDs (asterisk in C2) were rare, only six have been detected among the neurons assessed, two each in adult monkey, in ferret, and in human. In all cases the axon immediately bended down towards the white matter. Axon origins are marked by large arrows, small arrows indicate the course of biocytin-labeled axons. Scale bars 25 μm. The online version of this article includes the following figure supplement for figure 2: ***Figure 2–figure supplementmovie 1.*** The movie shows an AcD pyramidal neuron from infragranular layers of adult monkey cortex (center) and flanking non-AcD neurons labeled with SMI-32/βIV-spectrin.

In gray matter of non-primates, 10-21% of the pyramidal neurons were AcD cells (***Figure 3A***). The individual variability is given in ***Figure 3–figure supplement tables 1, 2.*** AcDs in rodent hippocampus are described as being functionally privileged which may be mirrored by their spine density. However, analysis of rat and ferret biocytin-stained pyramidal cells revealed that neither the dendrite sharing a root with an axon nor the AcDs had spine densities differing systematically from those of non-AcDs (***Figure 4***).

**Figure 3.**
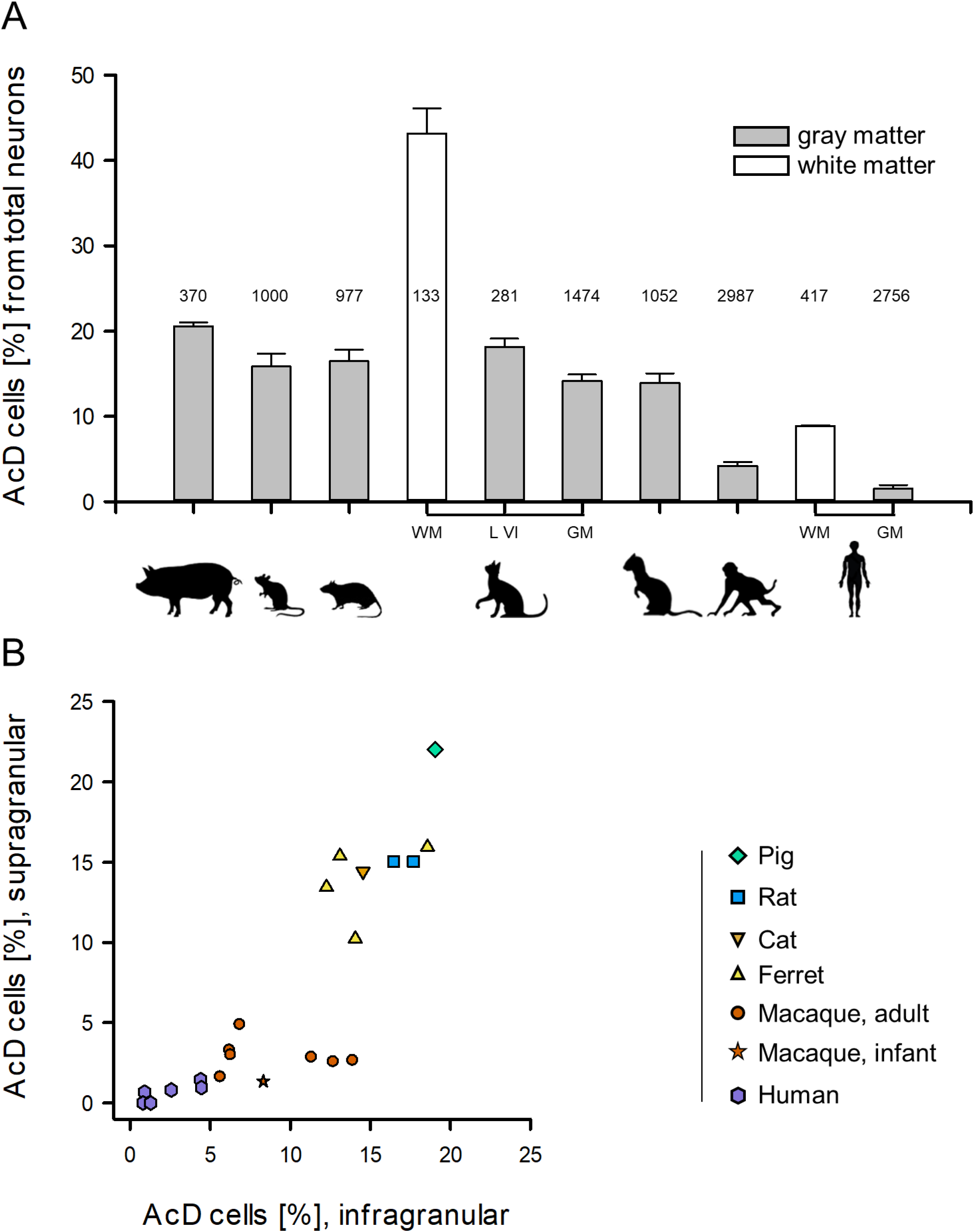
Proportion of AcD neurons across species. (**A**) Shown are mean ± S.E.M. of the individual percentages; values obtained by all staining methods employed per species have been pooled. Numbers in the bars are the total number of pyramidal neurons assessed. (**B**) Non-primate species showed roughly equal proportions of AcD neurons in supra- and infragranular layers. With some individual variability the range was 10-21%. In contrast, in monkey, the cluster was down-shifted due to overall much lower proportions. Further, it was right-shifted because supragranular pyramidal cells displayed much lower proportions of AcD cells compared to infragranular pyramidal cells. A Mann-Whitney rank sum test of “all primate” versus “all non-primate” proportions of supragranular and infragranular AcD cells, respectively, yielded p>0.001 for both comparisons. Comparing supragranular respectively infragranular values between macaque and human yielded p=0.001 and p<0.001, resp.. The online version of this article includes the following source data and figure supplement(s) for figure 1: ***Figure 3–figure supplement***

**Figure 3-figure supplement Table 1.** Proportion of pyramidal neurons with AcD: rodents. Individual proportions, number of cells assessed by the labeling methods indicated, and grand average is given, as well as individual laminar proportions. The latter have been obtained by a second round of quantification in order to control for reproducibility; in addition, sections have been assessed that had not been considered in the first round. Therefore, in Table 1, 2 and 3, the laminar percentages do not simply add up to proportions obtained for whole gray matter. Mouse Thy-1-EGFP expressing and βIV-spectrin+ pyramidal neurons vary from 10-22%, possibly due to the individual variability of the Thy-1 expression level. Thy-1 was only expressed in layer V and therefore, did not yield laminar percentages. n.a., not applicable.

|  |  |  |  |
| --- | --- | --- | --- |
| 240 | **Figure 3-figure supplement Table 1. Proportion of pyramidal neurons with AcD: rodents** |  |  |
|  | Species; cortical area;<br>staining method; age; sex | proportion of AcD cells [%];<br>n of cells assessed; | by layers:<br>supra %, infra % |
|  | **Mouse S1 cortex, layer V;**<br>**Thy1-EGFP/βIV-spectrin**<br>**immunofluorescence** |  |  |
|  | adult, female | 17.42 %, 178 cells | n.a. |
|  | adult, female | 11.78 %, 348 cells | n.a. |
|  | adult, female | 15.79 %, 36 cells | n.a. |
|  | adult, female | 18.84 %, 138 cells | n.a. |
|  | adult, male | 21.93 %, 187 cells | n.a. |
|  | adult, male | 14.82 %, 54 cells | n.a. |
|  | adult, male | 10.17 %, 59 cells | n.a. |
|  | *average [%], total n of cells* | **15.82 %, 1000 cells** |  |
|  | **Rat, visual cortex;**<br>**biocytin tracing** |  |  |
|  | adult, male, two hemispheres | 17.82 %, 174 cells | 15.06 %, 17.69 % |
|  | adult, male, two hemispheres | 15.07 %, 803 cells | 15.06 %, 16.46 % |
|  | *average [%], total n of cells* | **16.45 %, 977 cells** |  |

**Figure 3-figure supplement Table 2.** Proportion of pyramidal neurons with AcD: ungulate, carnivores. Individual proportions, number of cells assessed by the labeling methods indicated, and grand average is given, as well as individual laminar proportions. n.a., not applicable because no labeling of other layers; n.d., not determined due to too faint and erratic staining of supragranular layers.

| Species; cortical area;<br>staining method; age; sex | proportion of AcD cells [%];<br>n of cells assessed; | by layers:<br>supra %, infra % |
| --- | --- | --- |
| **Kitten visual cortex gyral white matter;**<br>**intracellular Lucifer Yellow** |  |  |
| P1, P2 (n = 2, sex n.d.) | 37.93 %, 58 cells | n.a. |
| P10, P11 (n = 2, sex n.d.) | 44.64 %, 56 cells | n.a. |
| P12, P14 (n = 2, sex n.d.) | 47.37 %, 19 cells | n.a. |
| *average [%], total n of cells* | **43.31 %, 133 cells** |  |
| **Kitten and adult cat visual cortex layer VI;**<br>**intracellular Lucifer Yellow** |  |  |
| P1, P5 (n = 2, sex n.d.) | 20.18 %, 114 cells | n.a. |
| P11 (n = 2, sex n.d.) | 17.65 %, 51 cells | n.a. |
| P19, P30 (n = 2, sex n.d.) | 14.89 %, 47 cells | n.a. |
| P52, P60 (n = 2, sex n.d.) | 20.00 %, 30 cells | n.a. |
| adult (n = 2, sex n.d.) | 17.95 %, 39 cells | n.a. |
| *average [%], total n of cells* | **18.13 %, 281 cells** |  |
| **Adult cat visual cortex;**<br>**immunofluorescence** |  |  |
| 3 months, sex n.d. | 13.35 %, 978 cells | 14.42 %, 14.55 % |
| adult, sex n.d. | 14.92 %, 496 cells | n.d. |
| *average [%], total n of cells* | **14.14 %, 1474 cells** |  |
| **Ferret, extrastriate cortex;**<br>**biocytin tracing** |  |  |
| adult, female | 16.56 %, 302 cells | 15.95 %, 18.57 % |
| adult, female | 14.66 %, 191 cells | 15.39 %, 13.11 % |
| adult, female | 11.30 %, 230 cells | 10.24 %, 14.06 % |
| adult, male | 13.07 %, 329 cells | 13.45 %, 12.26 % |
| *average [%], total n of cells* | **13.90 %, 1052 cells** |  |
| **Pig dorsoparietal cortex;**<br>**immunofluorescence** |  |  |
| 3 months, European wild boar, female | 20.11 %, 189 cells | 22.04 %, 19.05 % |
| 5 months, domestic, sex n.d. | 20.99 %, 181 cells | n.d. |
| *average [%], total n of cells* | **20.55 %, 370 cells** |  |

**Figure 3-figure supplement Table 3.** Proportion of pyramidal neurons with AcD: primates. Individual proportions, number of cells assessed by the labeling methods indicated, and grand average is given, as well as individual laminar proportions. Note the variable proportions of AcD neurons in infragranular layers and no obvious correlation between proportion and age of the individual macaques. Source data 1. Data from experiments plotted in *Figure 3A* Source data 1. Data and statistical analysis from experiments plotted in *Figure 3B.*

|  |  |  |  |
| --- | --- | --- | --- |
| 251 | **Figure 3-figure supplement Table 3. Proportion of pyramidal neurons with AcD: primates** |  |  |
|  | Species; cortical area;<br>staining method; age; sex | proportion of AcD cells [%];<br>n of cells assessed | by layers:<br>supra %, infra % |
|  | **Macaca mulatta, premotor cortex;**<br>**biocytin tracing** |  |  |
|  | Individual 1, 11 years, male | 4.93 %, 954 cells | 2.79 %, 11.38 % |
|  | Individual 2, 7 years, male | 3.10 %, 816 cells | 2.93 %, 6.32 % |
|  | Individual 3, 5 years, male | 3.80 %, 184 cells | 3.23 %, 6.25 % |
|  | **Macaca mulatta, parietal & visual cortex;**<br>**SMI-32/βIV-spectrin immunofluorescence** |  |  |
|  | Individual 4, 5 years, male | 5.86 %, 324 cells | 2.59 %, 13.95 % |
|  | Individual 5, 10 years, male | 5.20 %, 175 cells | 2.51 %, 12.72 % |
|  | **Macaca fascicularis, parietal cortex;**<br>**Golgi-Kopsch method** |  |  |
|  | Individual 6, adult, female | 3.58 %, 307 cells | 4.82 %, 6.90 % |
|  | Individual 7, adult, male | 3.07 %, 228 cells | 1.57 %, 5.67 % |
|  | *average [%] individuals 1-7, total n of cells* | **4.22 %, 2987 cells** | **2.92 %, 9.03 %** |
|  | **Macaca mulatta, visual cortex;**<br>**biocytin tracing** |  |  |
|  | Individual 8, P60, female | 4.51 %, 377 cells | 1.28 %, 7.85 % |
|  | **Macaca mulatta, premotor/M2 cortex;**<br>**SMI-32/βIV-spectrin immunofluorescence** |  |  |
|  | Individual 8, P60, female | 5.64 %, 1249 cells | 1.44 %, 8.78 % |
|  | *average [%] individual 8, total n of cells* | **5.08 %, 1626 cells** | **1.36 %, 8.32 %** |
|  | **Human, temporal lobe;**<br>**Golgi-Cox method** |  |  |
|  | Individual 1, 53 years, male | 2.56 %, 646 cells | 1.45 %, 4.40 % |
|  | Individual 2, 56 years, male | 2.79 %, 825 cells | 0.96 %, 4.66 % |
|  | **Human, auditory cortex;**<br>**Golgi-Kopsch method** |  |  |
|  | Individual 3, 63 years, male | 0.79 %, 253 cells | 0.67 %, 0.91 % |
|  | Individual 4, 71 years, female | 1.92 %, 677 cells | 0.80 %, 2.58 % |
|  | Individual 5, 75 years, male | 0.47 %, 215 cells | 0.00 %, 0.80 % |
|  | Individual 6, 88 years, female | 0.71 %, 140 cells | 0.00 %, 1.28 % |
|  | *average [%] individuals 1-6, total n of cells* | **1.52 %, 2756 cells** | **0.65 %, 2.44 %** |
|  | **Human, auditory cortex gyral white matter;**<br>**Golgi-Kopsch method** |  |  |
|  | Individual 3, 63 years, male | 8.69 %, 115 cells, | n.a. |
|  | Individual 4, 71 years, female | 8.88 %, 135 cells, | n.a. |
|  | Individual 5, 75 years, male | 8.70 %, 69 cells, | n.a. |
|  | Individual 6, 88 years, female | 9.18 %, 98 cells, | n.a. |
|  | *average [%] individuals 3-6, total n of cells* | **8.86 %, 417 cells** |  |

**Figure 4.**
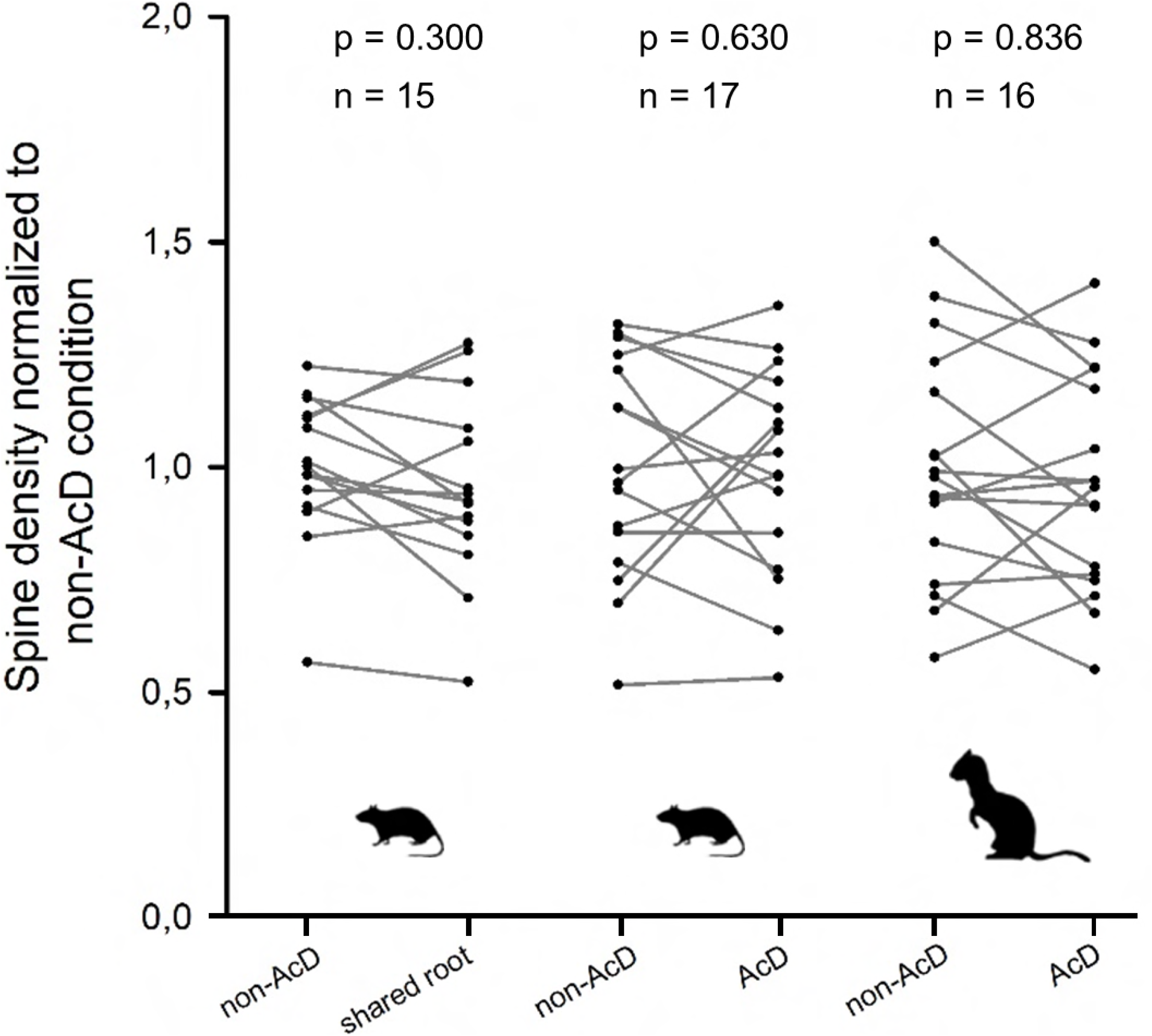
Spine density did not systematically differ between regular dendrites (non-AcDs), dendrites sharing a root with a neighboring axon, or AcDs. Data from adult rat and ferret biocytin material, values from each cell are connected by a line. For normalization, the average of the non-AcD has been set to 1, and all values were expressed relative to this. Mann-Whitney rank sum test p-values and the n are reported above each plot. Source data 2. Data and statistical analysis from experiments plotted in *Figure 4.*

In adult monkey gray matter only 3-6% of the pyramidal neurons were AcD cells. In human gray matter the proportion of AcD pyramidal neurons was 1.4% on average (***Figure 3A, Figure 3–figure supplement Table 3***). In the Golgi-Kopsch material, we assessed the proportion of “shared root” configuration. On average, an additional 6.17% of pyramidal cells of monkey gray matter (33 out of 535 cells, 4 individuals) and 3% of pyramidal cells of human gray matter (26 out of 875 cells, 4 individuals) had axons sharing a root with a basal dendrite.

A significant difference between non-primates and primates emerged after a layer-specific analysis. Non-primates had about equal proportions of AcD cells in supra- and infragranular layers. In contrast, monkey had 1-5% supragranular and 5-15% infragranular AcD cells; in human, the laminar percentages on average were 0.65% and 2.44%, respectively (***Figure 3B, Figure 3–figure supplement Table 3***). Primate and non-primate values differed significantly as did values of human and macaque (see ***Fig. 3B***, legend).

### Developmental aspects

Kitten layer VI pyramidal cells (***Figure 3A, Figure 3-figure supplement Table 2***) showed adult percentages of AcDs early postnatally. Many pyramidal cells were L-shaped or inverted-fusiform, with the axon emerging from one of the dominant dendrites (Lübke and Albus, 1989). In line with this, immunostained infant monkey M2/premotor cortex exhibited percentages of AcD cells comparable to adult monkey premotor cortex neurons labeled with biocytin, and again, AcD cells were more frequent in infragranular layers (***Figure 3A, Figure 3–figure supplement Table 3***).

Unexpectedly, of the pyramidal cells in kitten white matter (Wahle et al., 1994), 40.19% had axons emerging from the major dendrite (***Figure 3-figure supplement Table 2***). Even more striking, 8.86% of the interstitial pyramidal neurons of the human white matter (Meyer et al., 1992) displayed AcDs (***Figure 3–figure supplement Table 3***), and on average an additional 13.23% of the interstitial cells had axons emerging from a shared root. To summarize, we observed a substantial species difference with AcD cells being more frequent in non-primates. Within-species, we found clear laminar differences, with AcD cells being rare in primate supragranular layers, and more frequent in white matter (subplate/interstitial) neurons.

## Discussion

A clear majority of human gray matter pyramidal neurons have axons arising from the soma. In this aspect, in particular supragranular neurons of primates clearly differ from those of non-primates. Domestic pig and wild boar had similar proportions suggesting that domestication has no influence. Kitten and infant monkey data suggest that development has no impact on the overall proportions. Yet, in primates, ontogenetically older infragranular pyramidal cells display more AcDs than later-generated supragranular neurons. Neurons of the white matter seem to be a special case. In cat, they represent a subset of subplate cells transiently acting as instructors for the developing thalamocortical synaptic connectivity to layer IV (reviewed by Molnar et al., 2020). Pyramidal interstitial cells of adult primate may differ from the transient subplate of non-primate cortex (Meyer et al., 1992, Suarez-Sola et al., 2009; Sedmak and Judas, 2021). With regard to the functional concept of AcD neurons (Thome et al., 2014; Hamada et al., 2016; Kole and Brette, 2018), our findings suggest that action potential firing abilities bypassing somatic integration and inhibition are advantageous during development of thalamocortical wiring. White matter neurons reside at strategic positions to monitor incoming inputs and may quickly relay that information to the overlying gray matter. Assuming that subcortical afferents and white-to-gray matter projections match in topography (reviewed by Molnar et al., 2020), such a double-hit might narrow the time window of integration enabling synaptic plasticity or help to activate inhibitory neurons.

Electron microscopy has revealed that the AIS of axons originating from the apical dendrite of rat inverted pyramidal cells are shorter, thinner and less innervated by symmetric synapses than AIS of axons arising from the soma (Mendizabal-Zubiaga et al., 2007). In cat visual cortex, inverted-fusiform pyramidal neurons of layer VI serve corticocortical, but not corticothalamic projections; for instance, the feedback projection to area 17 from the suprasylvian sulcus (Einstein, 1996), an area involved in motion detection, processes optical flow and pupillary constriction. With regard to the functional concept of AcD cells, the kinetics of intra- and interareal information processing may have so far unrecognized facets.

Our data add to the view that human cortical pyramidal neurons differ in important aspects from those of non-primates (Elston et al., 2011, DeFelipe, 2011; Beaulieu-Laroche et al., 2018; Gidon et al., 2020; Rich et al., 2021). For instance, human supragranular pyramidal neurons have highly complex basal dendrites, each being a unique computational unit (reviewed by Goriounova and Mansvelder, 2019). Another recently described feature is the unique design of the human cortical excitatory synapses having pools of synaptic vesicles, release sites, and active zones that are much larger compared to those in rodents (Molnar et al., 2016; Yakoubi et al., 2019). Large and efficient presynapses may reliably depolarize the target cell’s somatodendritic compartment such that electrical dendroaxonic short circuits might become obsolete. We propose an evolutionary trend towards inputs that are conventionally integrated within the somatodendritic compartment and precisely modulated by inhibition to generate an optimally tuned behavioral output.

## Material and Methods

### Key resources table

| Reagent type | Species, label | Source or reference | Identifiers | Dilution used |
| --- | --- | --- | --- | --- |
| Antibody | anti-βIV-spectrin, rabbit | Höflin et al., 2017 | self-made | 1:500 |
| Antibody | anti-SMI-32 unphosphorylated neurofilaments, mouse | Covance, Muenster, Germany | Cat# SMI-32r, RRID: AB_2315331 | 1:1000 |
| Antibody | anti-NeuN, mouse | Merck (Millipore), Darmstadt, Germany | Cat # MAB377, RRID: AB_2298772 | 1: 1000 |
| Antibody | donkey anti-rabbit, Alexa-488 | Thermo Scientific, Waltham MA, USA | RRID: AB_2687506 | 1:1000 |
| Antibody | goat anti-rabbit, biotin | Dako A/S, Glostrup, Denmark | RRID: AB_2313609 | 1:1000 |
| Antibody | goat, anti-mouse Alexa-568 | Invitrogen, Carlsbad, CA, USA | RRID: AB_2534013 | 1:1000 |
| Antibody | sheep anti-mouse, biocytin | GE Healthcare Life Sciences, Braunschweig Germany, | Amersham Cat # RPN1001 | 1:200 |
| Chemical compound | Streptavidin Alexa-488 | Thermo Scientific, Waltham MA, USA | Cat #S11223 | 1:1000 |
| Chemical compound | ABC Elite horseradish peroxidase | Vector Labs Inc., Burlingame, CA, USA, | RRID: AB_2336827 | 1:250 |

### Animals

The data presented here were compiled by tissue sharing (immunohistochemistry) and from tissue that had originally been processed for unrelated projects, i.e. no additional animals were sacrificed specifically for this study.

### Biocytin injections

Two adult male rats received local biocytin injections into areas 17 and 18 in the course of teaching experiments done in the 1990s demonstrating surgery, tracer injections and biocytin histology. The animals were from the in-house breeding facility. The histological material has been used for decades to train neuroanatomy course students at the Department of Zoology and Neurobiology. Four adult pigmented ferrets (*Mustela putorius furo*) received BDA injections into the motion-sensitive posterior suprasylvian area (Philipp et al. 2005, Kalberlah et al. 2008). After a survival time of 6-13 days the animals were sacrificed and processed as described for the macaque cases. Three male adult macaque monkeys (*Macaca mulatta*) received tracer injections (15-20% biotin dextrane amine (BDA) MW 3000) into dorsal premotor cortex (Distler and Hoffmann, 2015). After a survival period of 14-17 days the animals were sacrificed with an overdose of pentobarbital and perfused through the heart with 0.9% NaCl and 1% procainhydrochloride followed by paraformaldehyde-lysine-periodate containing 4% paraformaldehyde in 0.1 M phosphate buffer pH 7.4. Coronal 50 μm thick frozen sections were cut on a microtome and processed for biocytin histochemistry with the avidin-biotin method (ABC Elite) with diaminobenzidine as chromogen enhanced with ammonium nickel sulfate (Distler and Hoffmann 2015). A P60 infant macaque received a biocytin injection into visual cortex and has been processed as above (Distler and Hoffmann, 2011).

### Intracellular Lucifer Yellow injections

The cat material was from a study on development of area 17 layer VI pyramidal cell dendrites (Lübke and Albus 1989). Briefly, Lucifer Yellow was iontophoretically injected into the somata in fixed vibratome slices of 100-150 μm thickness followed by UV-light photoconversion in the presence of diaminobenzidine towards a solid dark-brown reaction product. Further, we assessed neurons in the white matter of developing cat visual cortex prelabeled with the antibody “Subplate-1” followed by Lucifer Yellow injection and photoconversion (Wahle et al., 1994).

### Immunofluorescence

Adult macaque and P60 infant macaque material not used for immediate histological assessment had been stored after fixation and glycerol infiltration in isopentane at −80°C. From such spared blocks, 50 μm cryostat sections were cut for immunostaining. Adult cat material was from studies on development of visual cortex interneurons (Wahle and Meyer, 1987; Meyer and Wahle 1988). Cryoprotected slabs of these brains had been stored since then embedded in TissueTek at −80°C. The 5 month pig material was obtained from the Institutes of Physiology and Anatomy, Medical Faculty, University Mannheim (donated by Prof. Martin Schmelz). The P90 European wild boar material has been from current studies (Ernst et al., 2018; Sobierajski et al., 2021). Sections were pretreated with 3% H_2_O_2_ in TBS for 30 min, rinsed, incubated for 1 h in 0.5% Triton in TBS, blocked in 5% horse serum in TBS for 2 h followed by incubation in mouse anti-SMI-32 to stain somata and dendrites of subsets of pyramidal cells, and rabbit anti-βIV-spectrin (Höfflin et al., 2017) to stain the AIS. We could do only do 1 pig and 1 cat for the laminar analysis because the immunofluorescence did not to deliver sufficient basal dendritic SMI-32 labeling of supragranular neurons in the second available individual. Thus, for these two cases no reliable laminar data could be obtained. Mouse anti-NeuN staining of adjacent sections helped to identify the layers. After 48 h incubation at 8°C sections were rinsed, incubated in fluorescent secondaries including DAPI to label nuclei, and coverslipped for confocal analysis. Mouse material was collected as part of the ongoing dissertations of Nadja Lehmann and Susanna Weber, Institute of Neuroanatomy, Medical Faculty Mannheim, Heidelberg University, supervised by Prof. Maren Engelhardt. Sections were processed as described previously (Jamann et al. 2021). The intrinsic EGFP-signal was combined with βIV-spectrin immunostaining.

### Golgi impregnation

The Golgi-Cox impregnations were done with access tissue removed during transcortical amygdalo-hippocampectomy from two adult patients who suffered from temporal lobe epilepsy (individuals 1 and 2 in ***Figure 3-figure supplement Table 3***). All experimental procedures were approved by the Ethical Committees as reported (Schmuhl-Giesen et al., 2021, Yakoubi et al., 2019). These and other previous studies have demonstrated that the access tissue is normal because is is far from the epileptic focus. Biopsy tissue was processed using the Hito Golgi-Cox Optim-Stain kit (Hitobiotec Corp) as described (Schmuhl-Giesen et al., 2021; Yakoubi et al., 2019). Coronal sections (quality as shown by Schmuhl-Giesen et al. in their Figure 1-figure supplement 1) were analyzed for AcD neurons in supragranular (I-IV) and infragranular (V-VI) layers.

The Golgi-Kopsch impregnations of macaque cortex were done on spare tissue from experiments done by Prof. Dr. Barry B. Lee (Lee et al., 1983). The sections had been used as reference material in the Dept. of Anatomy, University of La Laguna, Tenerife, Spain. The Golgi-Kopsch impregnations of human material (individuals 3-6 in ***Figure 3-figure supplement Table 3***) was processed decades ago (Meyer 1987; Meyer at al., 1989; Meyer et al., 1992). The brains were from notarized donations to the Department of Anatomy of the University of La Laguna for teaching of medical students and for research. Donors had no neurological disorders. After death, the bodies were transferred to the Department and perfused with formalin. The brains were extracted, stored in formalin, and small selected blocks were processed using a variant of the Golgi-Kopsch method. Tissue blocks were immersed in a solution of 3.5 % potassium dichromate, 1% chloral hydrate, and 3% formalin in destilled water for 5 days, followed by immersion for 2 days in 0.75% silver nitrate. Blocks containing the auditory cortex (Heschl’s gyrus) were cut by hand with a razor blade, dehydrated and mounted in Epon. For the assessment of AcD cells in the white matter, the border between gray and white matter was traced (Meyer et al., 1992). We avoided this zone and took as orientation the dense aggregations of astrocytes in the white matter and the linear arrangement of blood vessels. As shown before (Meyer et al., 1992), interstitial pyramidal cells have a variety of shapes, from elongated bipolar to multipolar, but carry dendritic spines in contrast to non-pyramidal interstitial cells.

### Assignment of AcD

We assessed all pyramidal cells with sufficiently well stained basal and apical dendrites that had a recognizable axon. We analyzed fields of view where the labeled pyramidal cells are fairly perpendicularly oriented such that the apical dendritic trunk and the descending axon could be clearly seen. In the biocytin, Lucifer Yellow, and Golgi cases, the axon could be clearly distinguished from sometimes equally thin descending dendrites because the latter had spines. Moreover, the axons had a clear axon hillock, which is more prominent in primate > carnivore > rodent. Descending primary axons often gave rise to thinner collaterals. Note the complementary nature of the methods: Golgi impregnation labels neurons in all layers, with the Golgi-Cox method yielding a higher density of neurons than the Golgi-Kopsch method. Intracortical biocytin injections labels preferentially neurons with horizontal projections in layers II/III. SMI-32/βIV-spectrin labeling is strongest in infragranular layers, in particular layer V and large pyramidal cells of layer III.

All neurons fulfilling the criteria were sampled by 5 observers trained on the AcD criteria (M.E., Linz; G.M., La Laguna; P.W. together with I.G. or E.S., Bochum). View fields were arbitrarily selected. For light microscopy, neurons were viewed and scored with 40x and 63x objectives. For SMI-32/βIV-spectrin and Thy-1/βIV-spectrin fluorescence, images and the tile scan were done with a Leica TSC SP5 confocal microscope (40x and 10× objective resp., with 1.1 NA, 1024 × 1024 px). All AcD and non-AcD cells with sufficient staining of the initial dendrites were manually marked in the confocal stacks using the “3D-environment”-function of Neurolucida 360 similar to ***Figure 2-figure supplement movie 1*** (mp4) exported from the Leica programme. Global whole picture contrast, brightness, color intensity and saturation settings were adjusted with Adobe Photoshop^®^. Scale bars were generated with ImageJ (MacBiophotonics) and inserted with Adobe Photoshop^®^ (CS6 Extended, Version 13.0 x64).

The assignment was done in a very conservative manner accepting as AcD cells only neurons where the axon arose with recognizable distance to the soma or emerged at such an angle that a vector through the axon hillock will not project into the soma but bypasses the soma tangentially. Sometimes the axon and a dendrite emerged very close to each other or from a shared a root. Such cells have been accepted in recent studies already as AcD neurons (Hamada et al., 2016; Thome et al., 2014). We scored such cells as “somatic axon cells”. In some cases, we additionally scored this “shared root” configuration and percentages are given in the text. In particular the white matter pyramidal neurons of the human brain were difficult due to their elongated shape and the somata tapering into the major dendrites (Meyer et al., 1992). Therefore, we aimed for the clear-cut cases.

### Spine analysis

To elucidate if the privileged AcD has a higher spine density than non-AcD, spines were plotted with the Neurolucida (MicroBrightField Inc., Williston, VT, United States) at 1000x magnification from biocytin-labeled neurons of rat and ferret cortex from primary and secondary basal dendrites starting minimum 50 μm away from the soma. On average we were able to reconstruct 170 μm/neuron in rat and 145 μm/neuron in ferret. The number of spines per 100 μm dendritic length was computed and the value for the AcD was paired to the average value of the basal non-AcD of every neuron. Yet, the number of measurable neurons was limited for the following reasons. First, neurons had to be well backfilled with the tracer. Second, neurons had to have an appreciable length of the AcD plus a minimum of one basal non-AcD in the 50 μm thin sections. Third, these dendrites had to display branch orders of 2-4 because the proximally thicker stems are not suitable for spine analysis and often void of spines (Hübener et al., 1990). Fourth, only solitary cells residing not too close to the injection site with its high background could be analyzed. Spine densities varied in our data set. Technically, the degree of biocytin labeling expectedly varied with the strength of the connection to the injection site. Biologically, pyramidal cell type-specific spine densities are known to vary up to an almost spine-free state e.g. in Meynert cells (Hübener et al., 1990). To collect a sufficient n, we included moderately biocytin-backfilled cells, although they tended to present with a lower spine density. Moreover, most counts were taken from branch order 2-4 segments which may have less spines than terminal segments. Our density average in rat matched values reported for nonterminal segments of Golgi-stained near-adult hooded rat visual cortex supragranular pyramidal cells (Juraska, 1982). Our ferret spine values were lower compared to earlier reports (Clemo and Meredith, 2012) presumably for the reasons mentioned above. However, this would not compromize our finding because we compared only dendrites within the individual neurons. Would there be some systematic change of the spine density between the AcD and the non-AcD of each cell, the difference should manifest irrespective of the individual staining intensity.

### Analysis

Graphs and statistics were done with SigmaStat12.3 (Systat Software GmbH, Frankfurt am Main, Germany). We aimed at minimum 5 individual per group in order to run non-parametric Mann-Whitney rank sum tests. Source data for the graphs were included as excel files.

## Acknowledgements

PW and GM dedicate the paper to our friend and mentor Prof. Dr. Klaus Albus, who graciously declined to join as a coauthor although the cat material we investigated had been prepared in his lab at the Max-Planck-Institut für Biophysikalische Chemie, Göttingen, Germany. We thank Prof. Barry B. Lee, at that time at the Max-Planck-Institut für Biophysikalische Chemie, Göttingen, Germany, for sharing monkey brain material. We thank Prof. Dr. Klaus-Peter Hoffmann, Ruhr-Universität, Bochum, Germany, who led the studies delivering the material of rat, ferret and monkey. We thank Dr. Astrid Rollenhagen, JARA-Institute Brain Structure Function Relationship, Jülich, for advice with the human patient material.

## Data availability

All data obtained during the study are reported in the Figures and the Supplemental Tables.

## Additional Information

### Funding

Deutsche Forschungsgemeinschaft WA 541/13-1 and WA 541/15-1 to Petra Wahle. Deutsche Forschungsgemeinschaft EN 1240/2-1 to Maren Engelhardt. Deutsche Forschungsgemeinschaft Ho-450/25-1 and Deutsche Forschungsgemeinschaft SFB 509/A11 to Claudia Distler. The funding agency had no role in study design, data collection and interpretation, or the decision to submit the work for publication.

### Author contributions

Petra Wahle, Conceptualization, Methodology, Resources (cat and pig material); Investigation, Analysis, Visualization, Supervision, Writing - original draft, Writing – review and editing, funding acquisition;

Eric Sobierajski, Investigation, Analysis, Visualization;

Ina Gasterstädt, Investigation, Analysis, Visualization;

Susanna Weber, Investigation;

Nadja Lehmann, Investigation;

Joachim HR Lübke, Resources (cat and human material);

Maren Engelhardt, Resources (mouse material), Investigation, Analysis, Writing - original draft,

Writing – review and editing; funding acquisition;

Claudia Distler, Resources (monkey, ferret, rat material), Investigation, Writing - original draft,

Writing – review and editing, funding acquisition;

Gundela Meyer, Resources (human and monkey material), Investigation, Analysis, Writing - original draft, Writing – review and editing.

### Ethics

The data presented in this paper were collected via tissue sharing and from material that had originally been processed for projects not related to the present topic, i.e. no animals were sacrificed specifically for the present study.

## Notes

### Competing Interest Statement

The authors have declared no competing interest.

